# Development of High-Throughput Screening Assays for Inhibitors of ETS Transcription Factors

**DOI:** 10.1101/181420

**Authors:** Simon L. Currie, Stephen L. Warner, Hariprasad Vankayalapati, Xiao-Hui Liu, Sunil Sharma, David J. Bearss, Barbara J. Graves

## Abstract

ETS transcription factors from the ERG and ETV1/4/5 subfamilies are overexpressed in the majority of prostate cancer patients and contribute to disease progression. Here, we develop two *in vitro* assays for the interaction of ETS transcription factors with DNA that are amenable for high throughput screening. Using ETS1 as a model, these assays were applied to screen 110 compounds derived from a high-throughput virtual screen. We find that the use of lower affinity DNA-binding sequences, similar to those which ERG and ETV1 bind to in prostate cells, allowed for higher inhibition from many of these test compounds. Further pilot experiments demonstrated that the *in vitro* assays are robust for ERG, ETV1, and ETV5, three of the ETS transcription factors that are overexpressed in prostate cancer.

## Introduction

Site-specific transcription factors influence RNA polymerase activity in a gene-specific manner and are among the major factors that regulate normal development and define cellular fate. Transcription factors are often misregulated in human cancers, with the most abundant examples being the loss of the p53 tumor suppressor and overexpression of the C-MYC oncoprotein.^1^ Therefore transcription factors are highly desirable therapeutic targets. With the exception of steroid hormone receptors, transcription factors are difficult therapeutic targets due to the lack of highly concave ligand-binding surfaces. Nevertheless, there are recent examples demonstrating successful modulation of transcription factor activity through the inhibition of protein-protein and protein-DNA interfaces.^2-5^

The ETS family of transcription factors contains 28 human genes that have a conserved ETS DNA-binding domain (Fig. 1A). Factors of the ERG (ERG, FLI1, FEV) and ETV1/4/5 (ETV1, ETV4, ETV5) subfamilies are involved in chromosomal rearrangements that result in the overexpression of one of these proteins in the majority of prostate cancer patients.^6^ Preclinical modeling of prostate cancer suggests that the overexpression of ERG, ETV1, or ETV4 contributes to further disease progression, indicating that these transcription factors are desirable therapeutic targets.^7-8^

**Figure 1.**
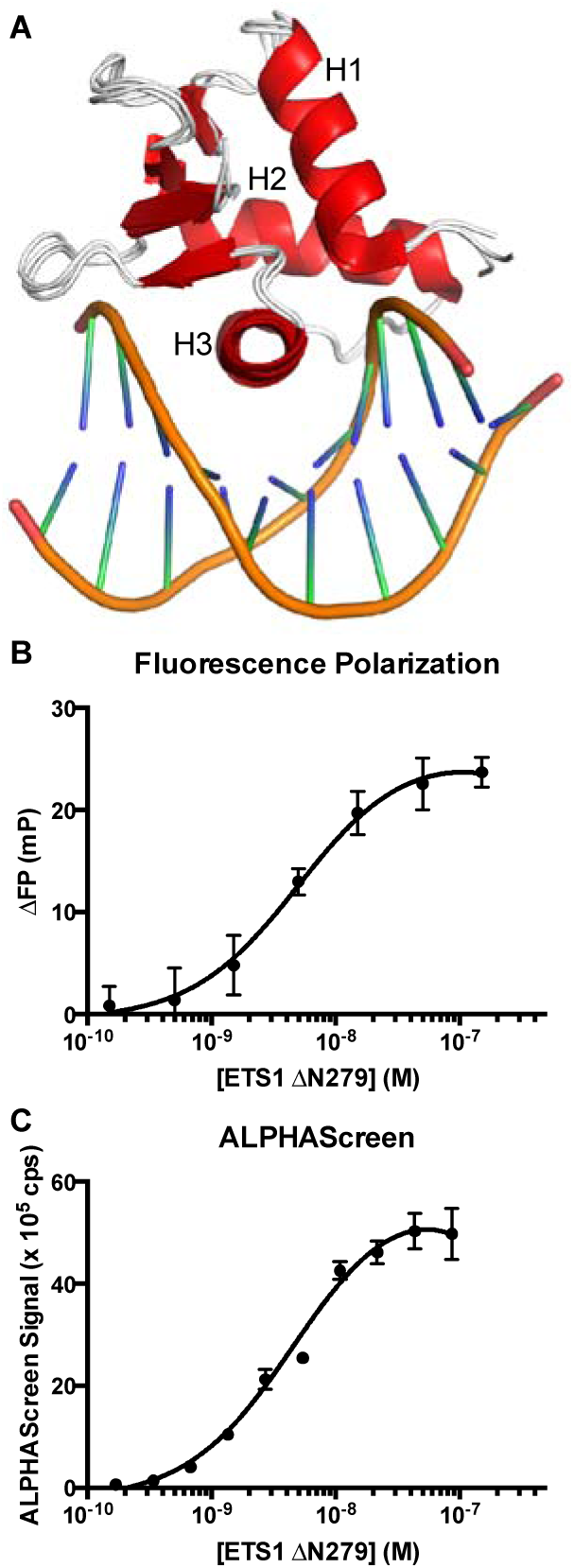
Assay development for ETS1-DNA interaction. (**A**) Structural alignment of ETS domains from ETS1 (PDB: 2NNY), ERG (4IRI), and ETV1 (4BNC) bound to DNA. H1, H2, and H3 indicate the order of theα-helices in the ETS domain from N-terminus to C-terminus. (**B**) Fluorescence polarization assay with a titration of ETS1 ΔN279 and 5 nM of 3’ fluorescein-labeled SC1 DNA. (**C**) ALPHAScreen assay with a titration of ETS1ΔN279 and 10 nM of 5’ biotin-labeled SC1 DNA.

Here we have designed *in-vitro* DNA-binding assays for ETS transcription factors that are amenable to high-throughput screening. We piloted these assays using ETS1 and a library of 110 compounds derived from high-throughput virtual screening. Furthermore, we demonstrate that using lower affinity ETS DNA binding sites, similar to those bound by ERG and ETV1 in prostate cancer cells, raises the efficacy of inhibitors of ETS-DNA interactions. Finally, we establish that these *in vitro* assays can be used with the prostate cancer relevant ETS factors ERG, ETV1, and ETV5.

## Materials and Methods

### DNA Constructs

Human cDNAs corresponding to full-length ETV1, ETV5, and ERG were cloned into the bacterial expression vector pET28 (Novagen) using standard sequence and ligation independent cloning strategies as previously described.^9^ ETS1 ΔN279 construct in pET28 was cloned as previously described.^10^

### Expression and Purification of Proteins

All proteins were produced in *Escherichia coli* (λDE3) cells. ETS1 ΔN279 efficiently expressed into the soluble fraction. Cultures of 1 L Luria broth (LB) were grown at 37 °C to OD_600_ ∼ 0.7 - 0.9, induced with 1 mM isopropyl-β-D-thiogalactopyranoside (IPTG), and grown at 30 °C for ∼ 3 hours.

Harvested cells were resuspended in 25 mM Tris pH 7.9, 1 M NaCl, 5 mM imidazole, 0.1 mM ethylenediaminetetraacetic acid (EDTA), 2 mM 2-mercaptoethanol (βME), and 1 mM phenylmethanesulfonylfluoride (PMSF). Cells were lysed by sonication and centrifuged at 40k rpm in a Ti-45 rotor (Beckmann) for at least 30 minutes at 4 °C. After centrifugation, the soluble supernatants were loaded onto a Ni^2+^ affinity column (GE Biosciences) and eluted over a 5-500 mM imidazole gradient. Fractions containing purified protein were pooled and dialyzed overnight at 4 °C into 25 mM Tris pH 7.9, 10% glycerol (v:v), 1mM EDTA, 50 mM KCl, and 1 mM dithiothreitol (DTT). After centrifugation at 40k rpm (Ti-45 rotor) for 30 minutes at 4 °C, the soluble fraction was loaded onto a SP sepharose cation exchange column (GE Biosciences) and eluted over a 50-1000 mM KCl gradient. Fractions containing the ETS proteins were loaded onto a Superdex 75 gel filtration column (GE Biosciences) and eluted fractions were analyzed by SDS-PAGE for purified ETS proteins. The final, purified protein was then concentrated on a 10-kDa molecular weight cut-off (MWCO) Centricon device, snap-frozen in liquid nitrogen, and stored at −80°C in single-use aliquots for subsequent *in vitro* studies.

Full-length ERG, ETV1, and ETV5 generally expressed more efficiently in the insoluble fraction using IPTG induction as described above. Harvested cells were resuspended as described above, sonicated and centrifuged at 15k rpm for 15 min at 4 °C. The soluble fraction was discarded and this procedure was repeated with the pellet / insoluble fraction twice more to rinse the inclusion bodies. The final insoluble fraction was resuspended with 25 mM Tris pH 7.9, 1 M NaCl, 0.1 mM EDTA, 5 mM imidazole, 2 mM BME, 1 mM PMSF, and 6 M urea. After sonication and incubation for ∼ 1 hr at 4 °C, the sample was centrifuged for 40k rpm for at least 30 min at 4 °C. The soluble fraction was loaded onto a Ni^2+^ NTA affinity column (GE Biosciences) and refolded by immediately switching to a buffer with the same components as above except lacking urea. After elution with 5 to 500 mM imidazole, the remaining purification steps, ion-exchange and size-exclusion chromatography, were performed as described above. However, a Q sepharose anion-exchange column was used instead of a SP sepharose cation-exchange column due to differing isoelectric points of the full-length proteins compared to ETS1 ΔN279.

Protein concentrations were measured using averages from the following two methods after ensuring that the concentrations from each method were in agreement with one another (within ∼ 2 fold). (1) Protein concentrations were determined by measuring the absorbance at 595 nm of 20 uL of protein combined with 1 mL of Protein Assay Dye Reagent (diluted 1:5 in deionized water)(Bio-Rad) and comparing to a bovine serum albumin standard curve. Molecular weights for each ETS protein were calculated using the Peptide Property Calculator (Northwestern). (2) Additionally, absorbance at a wavelength of 280 nm was measured on samples of protein mixed with 6 M Guanidine HCl (Thermo Scientific) at a 1:1 ratio and compared to a blank. Protein concentrations were determined using Beer’s Law (Abs280nm = ε\**l*\*c) with extinction coefficients for each protein calculated using Peptide Property Calculator (Northwestern).

### Electrophoretic Mobility Shift Assays

DNA-binding assays of ETS factors utilized duplexed 27-bp oligonucleotides with one of following two ETS sites. We first used the high-affinity consensus ETS binding site SC1 (Selected Clone 1): 5’-TCGACGGCCAAGCC**GGAA**GTGAGTGCC-3’ and 5’-TCGAGGCACTCAC**TTCC**GGCTTGGCCG-3’.^11^ Later optimizations used the lower-affinity ETS binding site SC13 (Selected Clone 13): 5’-TCGACGGCCAA<u>A</u>C<u>A</u>**GGA<u>T</u>**<u>A</u>T<u>C</u>AGTGCC-3’ and 5’-TCGAGGCACT<u>G</u>A<u>T</u>**<u>A</u>TCC**<u>T</u>G<u>T</u>TTGGCCG-3’.^11^ Boldface GGA(A/T) and (A/T)TCC indicate the core ETS binding site motif and underlined characters in SC13 indicate a difference compared to SC1. 0.2 nanomoles of each of these oligonucleotides, as measured by absorbance at 260 nM on a NanoDrop 1000 (Thermo Scientific), were labeled with [γ-^32^P] ATP (Perkin Elmer) using T4 polynucleotide kinase (Thermo Scientific) at 37° C for ∼ 30-60 min. After purification over a Bio-Spin® 6 chromatography column (Bio-Rad), the combined oligonucleotides were incubated at 100 °C for ∼ 5 min, and then cooled to room temperature over 1-2 hr. For binding reactions, the DNA concentration was diluted to 1 x 10^-11^ M and held constant, whereas protein concentrations ranged ∼ 6 orders of magnitude with the exact concentration range dependent on the equilibrium dissociation constant (*K*_*D*_) of the particular protein fragment. Protein concentration was determined after thawing each aliquot of protein, as described above. The binding reactions were incubated for 3 hr at 4° C in a buffer containing 25 mM Tris pH 7.9, 0.1 mM EDTA, 60 mM KCl, 6 mM MgCl_2_, 200 g/mL BSA, 10 mM DTT, 2.5 ng/ L poly(dIdC), and 10% (v:v) glycerol and then resolved on an 8% (w:v) native polyacrylamide gel at 4 °C. The ^32^P-labeled DNA was quantified on dried gels by phosphorimaging on a Typhoon Trio Variable Mode Imager (Amersham Biosciences). *K*_*D*_ values were determined by nonlinear least squares fitting of the total protein concentration [P]_t_ versus the fraction of DNA bound ([PD]/[D]_t_) to equation (1) using Kaleidagraph (v. 3.51; SynergySoftware). Due to the low concentration of total DNA, [D]_t_, in all reactions, the total protein concentration is a valid approximation of the free, unbound protein concentration.

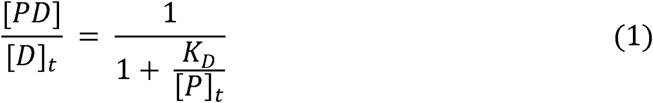

### Computational Methods

Computational methods were used as previously described.^12^ All computational studies used PDB ID 2NNY for the structural coordinates of ETS1.^13^ PocketFinder (ICM) and SiteMap (Schrödinger) were used to define ligand-binding sites. Out of the three ETS1 protein and one DNA ligand-binding sites that were defined by PocketFinder and SiteMap, only ETS1 site 1 was used for docking studies.

The compound database was prepared using Ligprep 2.1.23 of the Schrodinger Suite and ICM’s inbuilt preparation of three-dimensional ligands. A small molecule ligand library of 13 million compounds was docked against ETS1 using Glide High Throughput Virtual Screen. The top ∼ 15% ranked compounds were then redocked with the relatively more computationally expensive Glide standard precision scoring. The top ∼ 0.5% ranked were then subjected to further virtual screening using Glide extra-precision and ICM docking and scoring methods.

The final compounds that were identified for *in vitro* screening were the top ranking compounds from this final round of virtual screening that also met certain physicochemical criteria, such as solubility > 50 μg/mL, permeability > 50 nmol/s, and polar surface area < 120 Å^2^ as determined by QikProp. In addition to these rankings, redundant compounds were removed using ICM Molcart to improve the chemical diversity of the final set of compounds. Visual inspection of the docking results was used to evaluate binding mode, position, and orientation. In sum, this process resulted in 110 compounds that were purchased and screened using the *in vitro* ETS1-DNA binding assays.

### Fluorescence Polarization

Fluorescence polarization reactions were performed in the same buffer as described above for EMSAs. SC1 DNA was ordered with a 3’ fluorescein. Reactions were carried out in 20 μL volumes in black 384 well plates (Corning). The protein, DNA, and compound were incubated for 30 min at room temperature, protected from light. Timecourse studies demonstrated that less than 5 min were required for the protein-DNA reaction to reach equilibrium; however, we went with a longer incubation time to encourage compound – protein interactions, potentially with significantly lower affinity and kinetics, to also reach equilibrium. Reactions containing up to 5% DMSO showed no influence on the DNA-protein interaction. Plates were then analyzed on an Envision 2104 Multilabel Reader (Perkin Elmer). To calculate percent inhibition the signal (mp) for each compound was compared to positive (10 nM protein, 5 nM DNA, 0 μM compound; set to 0% inhibition) and negative (0 nM protein, 5 nM DNA, 0 μM compound; set to 100% inhibition) controls.

### ALPHAScreen

ALPHAScreen reactions were performed in the same buffer as described above for EMSAs except without 10% glycerol as this caused uneven distribution of the ALPHA beads. SC1 and SC13 DNA were ordered with a 5’ biotin. Reactions were carried out in 25 μL volumes in 384 well white OptiPlate-384 HB plates (Perkin Elmer). ALPHAScreen was performed according to manufacturer’s recommendations. Briefly, protein, compound, and DNA were incubated at room temperature for 60 min, protected from light. Nickel chelate acceptor beads were then added followed by another 60-min incubation at room temperature, protected from light. Then streptavidin donor beads were added followed by another 60-min incubation at room temperature. Plates were then analyzed on an Envision 2104 Multilabel Reader (Perkin Elmer). To calculate percent inhibition the signal (cps) for each compound was then compared to positive (10 nM protein, 10 nM DNA, 0 μM compound; set to 0% inhibition) and negative (0 nM protein, 10 nM DNA, 0 μM compound; set to 100% inhibition) controls.

### Comparison of Fluorescence Polarization and ALPHAScreen Assays

Equation (2) was used to compare the assay performance between fluorescence polarization and ALPHAScreen assays for ETS1 (μ and σ are mean and standard deviation and c+ and c- are positive and negative controls).^14^

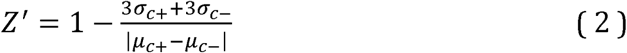

### Results and Discussion

ETS1 ΔN279 (residues 279 – 441) was used to pilot *in vitro* assays that could be utilized for high-throughput screening of potential small molecule inhibitors of ETS-DNA interactions. This fragment has robust expression in a recombinant system and contains the same affinity for DNA as full-length ETS1.^10^ The ETS domains of ETS1, ERG, and ETV1 are sequentially and structurally conserved (**Fig. 1A** and **Suppl. Fig. S1A**). Therefore, ETS1 serves as a good model for the DNA binding of these other ETS factors, and inhibitors that prevent ETS1 from binding to DNA would likely also inhibit ERG and ETV1.

ETS1 ΔN279 was expressed in *E. coli* and purified using a Ni^2+^ affinity column, a cation exchange column, and a size exclusion column (**Suppl. Fig. S1B**). Using <u>e</u>lectrophoretic <u>m</u>obility <u>s</u>hift <u>a</u>ssay (EMSA) we measured the binding of ETS1 ΔN279 to DNA with a consensus ETS site (5’-CCGGAAGT-3’), termed SC1 (<u>S</u>elected <u>C</u>lone 1).^11^ The measured *K*_*D*_ of 0.4 nM is in agreement with previous measurements for this fragment binding to DNA (**Suppl. Fig. S1C**).^10^ The yield of ETS1 ΔN279 was approximately five milligrams of purified protein per liter of bacterial culture, which provided plenty of protein for this study and could be efficiently scaled up to provide enough protein for a high-throughput *in-vitro* screen.

We next optimized screening conditions with the validated ETS1 ΔN279 for two potential high-throughput assays: fluorescence polarization, and ALPHAScreen. The fluorescence polarization assay uses a fluorescein-tagged SC1 DNA and measures the change in the polarization of fluorescently emitted light when the DNA is free in solution versus when the DNA is bound by a transcription factor. The ALPHAScreen assay brings beads that engage in fluorescence resonance energy transfer (FRET) into proximity though conjugation to a transcription factor and its recognition DNA site using Ni^2+^-His_6_ and streptavidin-biotin interactions, respectively. First, titration of DNA demonstrated that using 5 nM of fluorescein-tagged DNA for fluorescence polarization or 10 nM of biotin-tagged DNA for ALPHAScreen minimized the amount of DNA while still retaining a robust signal in these assays with ETS1 ΔN279. With these amounts of DNA, titration of ETS1 ΔN279 showed a dose-dependent response in these two assays with a concentration of around 30-50 nM generating maximum signal (**Fig. 1B,C**). Based on these titrations, 10 nm concentrations of ETS1 ΔN279 were used in the fluorescence polarization and ALPHAScreen assays for compound screening studies. The maximum signal and the baseline were used to calculate a Z’ factor for these assays. The fluorescence polarization assay had a Z’ factor of 0.4 and the ALPHAScreen assay had a Z’ factor of 0.7. Z’ factors above 0.5 are considered to be excellent assays for high-throughput screening purposes.^14^ Whereas the ETS1 ALPHAScreen assay already clears this guideline, the ETS1 fluorescence polarization assay is close and could likely be optimized to achieve Z’ factors over 0.5.

Computer modeling was utilized to enrich for likely bioactive compounds to be screened using these newly established *in-vitro* assays. Briefly, PocketFinder (ICM) and SiteMap (Schrodinger) were used to define ligand-binding pockets in the ETS domain of ETS1 (**Suppl. Fig. 2**). Sequential rounds of virtual screening using one of these defined ligand-binding pockets, termed ETS1 site 1, culled a starting library of 13 million compounds down to 110 compounds to be tested in the *in-vitro* ETS1 DNA binding assays. In addition to the predicted strength of interaction with ETS1, these compounds were also filtered to optimize chemical diversity and enrich for compounds with favorable physicochemical properties.

**Figure 2.**
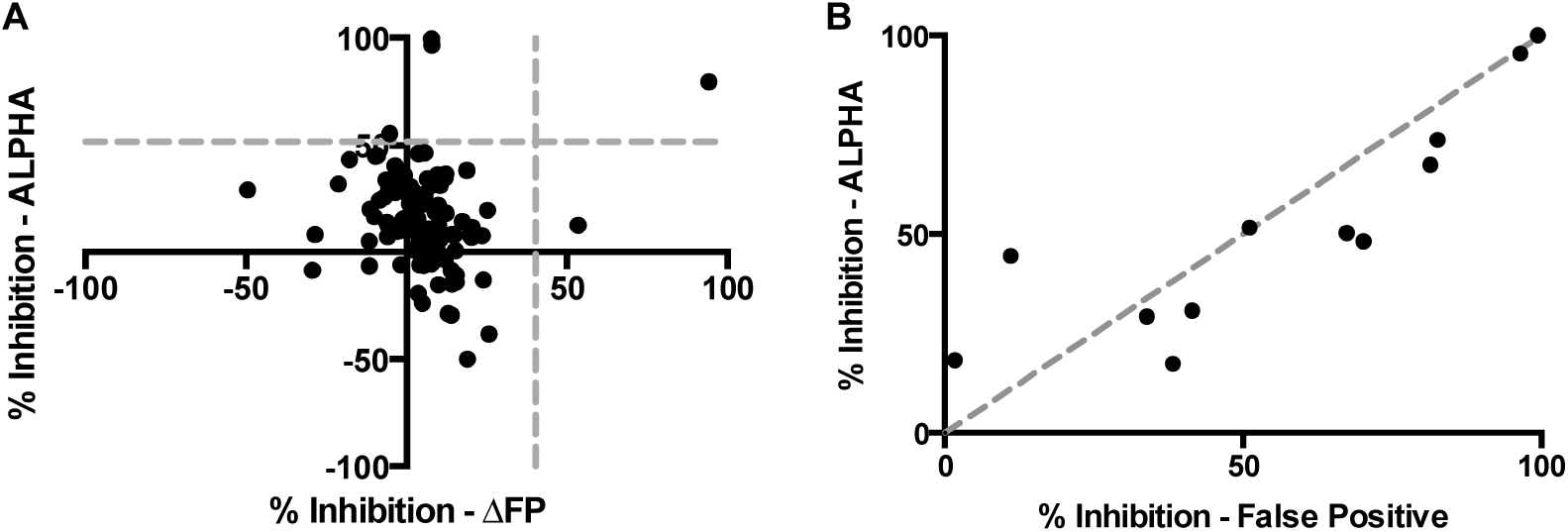
*In vitro* screens for inhibitors of ETS1–SC1 DNA interaction with fluorescence polarization and ALPHAScreen. (**A**) 110 compounds identified from virtual screening (Suppl. Fig. 2) were assayed for inhibition of ETS1-SC1 DNA interaction using fluorescence polarization (ΔFP) and ALPHAScreen (ALPHA). Percent inhibition was calculated with reference to positive (protein and DNA, no compound) and negative (DNA only, no protein or compound) controls. Dotted gray lines indicate three standard deviations separation from the baseline for each assay. (**B**) Counterscreen of the top hits from ALPHAScreen assay using the TruHits false positive kit. Dotted gray line indicates equivalent inhibition of the ALPHAScreen assay and the false positive assay.

A constant concentration of protein and DNA, as indicated above, was used to test the inhibition of each of the 110 compounds that resulted from virtual screening. These compounds were tested at a single concentration of 60 μM and each compound or control was measured in quadruplicates. Using three standard deviations above the baseline (3-SD) as a cutoff, only two compounds in the fluorescence polarization assay, and four compounds in the ALPHAScreen assay, respectively, inhibited the ETS1 ΔN279-DNA interaction. Only one of these compounds inhibited this interaction in both assays (**Fig. 2A**).

To further investigate these compounds, as well as some additional compounds that were close to the 3-SD cutoff, we utilized the ‘TruHits’ false positive screen in ALPHAScreen. In this assay a small molecule that covalently-conjugates biotin and His_6_ together is used in lieu of the biomolecules of interest, in this case ETS1 ΔN279 and SC1 DNA. Compounds that inhibit the false positive assay do so through a manner inherent to the assay itself such as by absorbing light in the donor or emission wavelengths or by disrupting the streptavidin-biotin or His_6_ - Ni^2+^ interactions that conjugate the biomolecules to the ALPHA beads. All of the compounds that strongly inhibited the ALPHAScreen assay also strongly inhibited this false positive assay (**Fig 2B**). Only two compounds that had weak to moderate inhibition of the ALPHAScreen assay displayed differential preference for inhibiting the ALPHAScreen assay more robustly than the false positive assay.

With very few, if any, actual hits from our first round of *in vitro* screening we next considered potential adjustments to our assays. One potential challenge with this screen is that the strength of the ETS1 ΔN279-SC1 DNA interaction (*K*_*D*_ = 0.4 nM) might conceal the discovery of starting compounds with relatively lower affinity for ETS1 ΔN279, which then could be further optimized for inhibition. To address this we switched from SC1 (5’-GCCGGAAGTG-3’), the highest affinity DNA sequence for ETS1, to a weaker ETS1 binding sequence, SC13 (5’-<u>A</u>C<u>A</u>GGA<u>TA</u>T<u>C</u>-3’). By EMSA, ETS1 ΔN279 bound to SC13 with *K*_*D*_ of 3 nM (data not shown). This roughly tenfold weaker interaction is consistent with the difference observed between SC1 and SC13 DNA with other ETS1 truncations.^11^

We rescreened the 110 compounds against ETS1 and SC13 DNA. Eighteen of these compounds inhibited the ETS1-SC13 interaction above the 3-SD cutoff, as compared to only four for the ETS1-SC1 interaction (**Fig. 3A**). While many of these compounds still inhibited the ‘TruHits’ false positive assay, 12/20 compounds showed more inhibition in the ETS1-SC13 assay than the false positive assay (**Fig. 3B**), compared to only 2/12 compounds that showed more inhibition in the ETS1-SC1 assay than the false positive assay (**Fig. 2B**). Therefore, using the weaker interaction of ETS1 with SC13 DNA enabled higher disrupt of the ETS1-DNA interaction by compounds. Additionally, a significant part of the inhibition observed in the ALPHAScreen assays for most of these compounds appears to come from “off-target” effects in the assay, besides interrupting the ETS-DNA interaction. However, as several of these compounds display stronger inhibition of the ETS1-SC13 assay than the false positive assay, these compounds may inhibit the ETS1-DNA interaction in addition to the ALPHAScreen assay in general.

**Figure 3.**
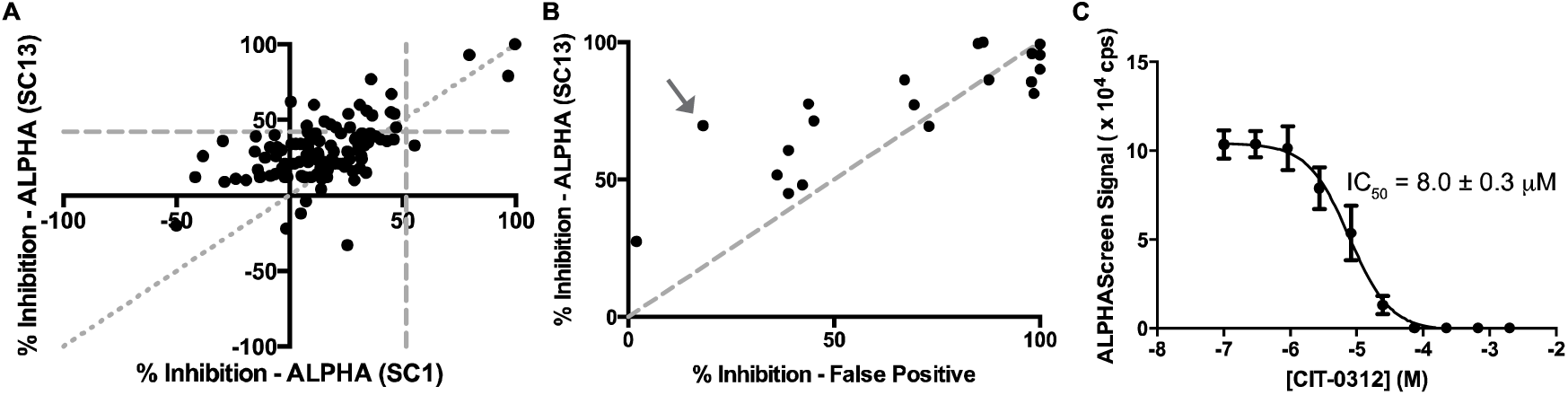
Screen for inhibitors of ETS1–SC13 DNA interaction using ALPHAscreen. (**A**) Comparison of compound inhibition against ETS1-SC1 DNA (x-axis) and ETS1-SC13 DNA (y-axis). SC13 is a lower-affinity ETS binding DNA sequence. The horizontal and vertical dotted gray lines indicate three standard deviations separation from the baseline for each screen. The diagonal, finely dotted gray line indicates equivalent inhibition against both of the DNA sequences. (**B**) Counterscreen of the top hits from ETS1-SC13 assay using the TruHits false positive kit. Dotted gray line indicates equivalent inhibition of the ALPHAScreen assay and the false positive assay. Arrow indicates the compound with the largest differential of inhibition of ETS-DNA assay compared to false positive assay that was used for further studies. (**C**) Representative titration of compound CIT-0312 using ALPHAScreen assay with ETS1 ΔN279 and SC13 DNA. Indicated IC_50_ of 8.0 ± 0.3 μM (mean ± standard deviation) for this compound was calculated from three replicate experiments.

In both the ETS1-SC1 and ETS1-SC13 screens the same compound, CIT-0312, displayed the largest differential between inhibition of ETS1 ΔN279-DNA assays and inhibition of the false positive assay. Therefore, this compound displayed the most specific inhibition of the ETS1-DNA interaction. Additionally, CIT-0312 more robustly inhibited the ETS-SC13 interaction (73%) than the ETS-SC1 interaction (36%), as would be expected given the relatively weaker affinity of the ETS-SC13 interaction. This compound inhibited ETS1 ΔN279-SC13 DNA interaction in the ALPHAScreen assay with an IC_50_ of 8.0 ± 0.3 μM (mean ± standard deviation). To confirm this inhibition we tested CIT-0312 using EMSAs. In this orthogonal assay CIT-0312 inhibited the ETS1 ΔN279-SC1 DNA with an IC_50_ of 27 ± 5 μM (**Suppl. Fig. 3**). Further investigation demonstrated that this compound lacked specificity as it similarly inhibited cJUN-FOS and FOXA1 transcription factors from binding to their cognate DNA recognition sites (Data not shown). Therefore, this particular compound must be inhibiting the DNA binding of ETS1, as well as other transcription factors, through a non-specific mechanism that is distinct from the prediction of our *in silico* modeling (**Suppl. Fig. 2**).

Within the ETS family of transcription factors, ERG and ETV1/4/5 subfamily proteins are overexpressed in prostate cancer and contribute to disease progression, making therapeutic inhibition of these proteins desirable.^7-8^ To establish that the screening assays used here for ETS1 are also suitable for these oncogenic proteins we expressed and purified full length, His_6_-tagged ERG, ETV1, and ETV5. Titrations of these proteins with 10 nM of biotin-tagged SC13 DNA and streptavidin donor and nickel chelate acceptor beads established that these proteins similarly generate robust ALPHAScreen signal, with a maximum signal observed around 20-70 nM, depending on the individual protein (**Fig. 4**). Each of these interactions had Z’ factors over 0.5 (ERG, 0.8; ETV1, 0.6; ETV5, 0.8) suggesting that they would be suitable for high-throughput screening.

**Figure 4.**
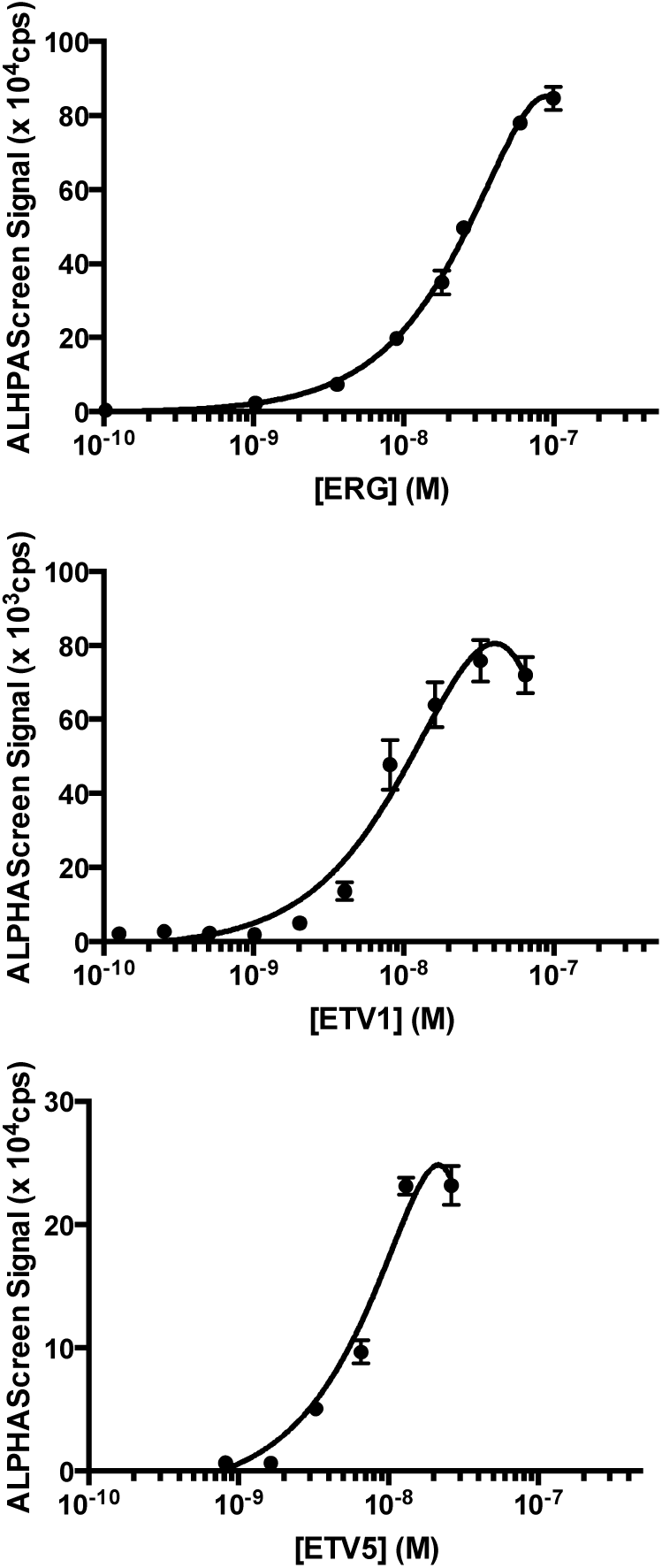
Assay development for ERG-, ETV1-, and ETV5-DNA interactions. ALPHAScreen assay with a titration of ERG (top), ETV1 (middle), or ETV5 (bottom) and 10 nM 5’ biotin-labeled SC13 DNA.

In summary we have established two *in vitro* assays, ALPHAScreen and fluorescence polarization, that are suitable for high-throughput screening of potential small molecule inhibitors of ETS–DNA interactions. Using weaker affinity DNA, such as SC13, was advantageous for more readily identifying potential lead compounds from the screens. Interestingly, these weaker affinity DNA sites may also be more biologically relevant as they more closely resemble the ERG and ETV1 DNA-binding sites that are relevant in prostate cancer.^7-8^ In contrast, consensus ETS sites, such as SC1, are redundantly regulated by multiple ETS factors and control the expression of housekeeping genes.^15^ These *in vitro* assays are suitable for ETS factors with high clinical relevance, such as ERG and ETV1, and can be used for performing high-throughput screens of these factors. Furthermore, as directly inhibiting transcription factor-DNA interactions remains a difficult target,^1^ these assays could also be used to screen for compounds that inhibit the function of ETS factors through alternative mechanisms. For instance, screens could conducted for small molecules that reinforce the diverse autoinhibitory mechanisms of ETS factors.^9^ Alternatively, these assays could be readily modified to search for disruptors of protein-protein interactions between ETS factors and important transcriptional coregulators.

## Acknowledgements

The authors are grateful to Charles Meeker, Christian Millsop, Katherine Chandler, and Krista Meyer for invaluable discussions, and Jack Skalicky, Frank Whitby, and Christopher Hill for advice and collaboration on structural experiments.

## References

1. Bhagwat, A. S.; Vakoc, C. R., Targeting transcription factors in cancer. Trends Cancer 2015, 1 (1), 53–65.

2. Illendula, A.; Pulikkan, J. A.; Zong, H., et al., Chemical biology. A small-molecule inhibitor of the aberrant transcription factor CBFbeta-SMMHC delays leukemia in mice. Science 2015, 347 (6223), 779–84.

3. Wang, X.; Qiao, Y.; Asangani, I. A., et al., Development of Peptidomimetic Inhibitors of the ERG Gene Fusion Product in Prostate Cancer. Cancer Cell 2017, 31 (4), 532–548 e7.

4. Butler, M. S.; Roshan-Moniri, M.; Hsing, M., et al., Discovery and characterization of small molecules targeting the DNA-binding ETS domain of ERG in prostate cancer. Oncotarget 2017.

5. Pop, M. S.; Stransky, N.; Garvie, C. W., et al., A small molecule that binds and inhibits the ETV1 transcription factor oncoprotein. Mol Cancer Ther 2014, 13 (6), 1492–502.

6. Robinson, D.; Van Allen, E. M.; Wu, Y. M., et al., Integrative clinical genomics of advanced prostate cancer. Cell 2015, 161 (5), 1215–28.

7. Baena, E.; Shao, Z.; Linn, D. E., et al., ETV1 directs androgen metabolism and confers aggressive prostate cancer in targeted mice and patients. Genes Dev 2013, 27 (6), 683–98.

8. Chen, Y.; Chi, P.; Rockowitz, S., et al., ETS factors reprogram the androgen receptor cistrome and prime prostate tumorigenesis in response to PTEN loss. Nat Med 2013, 19 (8), 1023–9.

9. Currie, S. L.; Lau, D. K. W.; Doane, J. J., et al., Structured and disordered regions cooperatively mediate DNA-binding autoinhibition of ETS factors ETV1, ETV4 and ETV5. Nucleic Acids Res 2017, 45 (5), 2223–2241.

10. Lee, G. M.; Pufall, M. A.; Meeker, C. A., et al., The affinity of Ets-1 for DNA is modulated by phosphorylation through transient interactions of an unstructured region. J Mol Biol 2008, 382 (4), 1014–30.

11. Nye, J. A.; Petersen, J. M.; Gunther, C. V., et al., Interaction of murine ets-1 with GGA-binding sites establishes the ETS domain as a new DNA-binding motif. Genes Dev 1992, 6 (6), 975–90.

12. Sorna, V.; Theisen, E. R.; Stephens, B., et al., High-throughput virtual screening identifies novel N'-(1-phenylethylidene)-benzohydrazides as potent, specific, and reversible LSD1 inhibitors. J Med Chem 2013, 56 (23), 9496–508.

13. Lamber, E. P.; Vanhille, L.; Textor, L. C., et al., Regulation of the transcription factor Ets-1 by DNA-mediated homo-dimerization. EMBO J 2008, 27 (14), 2006–17.

14. Zhang, J. H.; Chung, T. D.; Oldenburg, K. R., A simple statistical parameter for use in evaluation and validation of high throughput screening assays. J Biomol Screen 1999, 4 (2), 67–73.

15. Hollenhorst, P. C.; Chandler, K. J.; Poulsen, R. L., et al., DNA specificity determinants associate with distinct transcription factor functions. PLoS Genet 2009, 5 (12), e1000778.

